# Predicting long-term Type 2 Diabetes with Support Vector Machine using Oral Glucose Tolerance Test

**DOI:** 10.1101/688804

**Authors:** Hasan Abbas, Lejla Alic, Madhav Erraguntla, Jim Ji, Muhammad Abdul-Ghani, Qammer Abbasi, Marwa Qaraqe

**Affiliations:** Department of Electrical & Computer Engineering, Texas A&M University at Qatar, Doha 23874, Qatar.; Magnetic Detection & Imaging group, Faculty of Science & Technology, University of Twente, Enschede, The Netherlands.; Department of Industrial & Systems Engineering, Texas A&M University, College Station, TX 77843, USA.; Electronic & Nanoscale Engineering, University of Glasgow, Glasgow, Scotland G12 8QQ.; UT Health, San Antonio, TX 78229, USA.; College of Science and Engineering, Hamad Bin Khalifa University Doha, Qatar.

## Abstract

Diabetes is a large healthcare burden worldwide. There is substantial evidence that lifestyle modifications and drug intervention can prevent diabetes, therefore, an early identification of high risk individuals is important to design targeted prevention strategies. In this paper, we present an automatic tool that uses machine learning techniques to predict the development of type 2 diabetes mellitus (T2DM). Data generated from an oral glucose tolerance test (OGTT) was used to develop a predictive model based on the support vector machine (SVM). We trained and validated the models using the OGTT and demographic data of 1,492 healthy individuals collected during the San Antonio Heart Study. This study collected plasma glucose and insulin concentrations before glucose intake and at three time-points thereafter (30, 60 and 120 min). Furthermore, personal information such as age, ethnicity and body-mass index was also a part of the dataset. Using 11 oral glucose tolerance test (OGTT) measurements, we have deduced 61 features, which are then assigned a rank and the top ten features are shortlisted using Minimum Redundancy Maximum Relevance feature selection algorithm. All possible combinations of the 10 best ranked features were used to generate SVM based prediction models. This research shows that an individual’s plasma glucose levels, and the information derived therefrom have the strongest predictive performance for the future development of T2DM. Significantly, insulin and demographic features do not provide additional performance improvement for diabetes prediction. The results of this work identify the parsimonious clinical data needed to be collected for an efficient prediction of T2DM. Our approach shows an average accuracy of 96.80 % and a sensitivity of 80.09 % obtained on a holdout set.

## Introduction

The global incidence of diabetes was estimated at 422 million in the year 2014 and its prevalence among the adult population increased from 4.7 % in 1980 to 8.5 % in 2014 [1]. In 2015 alone, about 1.6 million deaths worldwide were attributed to diabetes. In addition to the high mortality rate, an individual with diabetes is at a greater risk of developing cardiovascular disease (CVD), visual impairment and limb amputations, as compared to a non-diabetic individual. Due to the substantial socio-economic burdens that are associated with diabetes, its early detection, prevention, and management has become a worldwide top-level health concern. There is experimental evidence that the development of diabetes can be delayed or even prevented provided an individual undertakes a lifestyle change that includes diet management, adopting exercise, and adhering to a pharmacological treatment [2]. The early identification of high risk individuals of diabetes is therefore, essential for targeted prevention strategies [3].

Even though the number of clinical studies aimed at diagnosing diabetes has been growing recently, studies predicting the risk of developing diabetes are limited. This subject has lately received an increased amount of research interest [4]. However, the clinical significance of such predictions largely depend on the type and quality of data collected. There are studies that assign a probability to the future risk of diabetes using socio-demographic characteristics such as age, ethnicity, body-mass index (BMI) and genealogical information collected through population [5,6]. Due to the unreliable data collection, such techniques can be misleading. The collection of blood samples, on the other hand, provides more reliable data and is a first step towards the disease prognosis with a deeper clinical insight [7]. The OGTT is commonly used to screen diabetes [8] and to provide a critical understanding of its future evolution [9]. In an OGTT, the plasma glucose and insulin levels are measured at regular intervals in a 2-hr period after orally administering a standard dose of glucose [9]. The glucose tolerance and insulin resistance are two of the most significant parameters deduced from the OGTT that are widely regarded as the major factors in the development of type 2 diabetes mellitus (T2DM).

A precursory stage of diabetes, commonly referred to as prediabetes, exists before overt T2DM, and is described by impaired fasting glucose (IFG), along with impaired glucose tolerance (IGT). According to the World Health Organization (WHO) diagnostic criteria, the IFG is defined as fasting plasma glucose level of 100 mg dL to 125 mg dL. The IGT which describes an abnormally raised glucose level is defined as the 2-hour plasma glucose level in the range of 140 mg dL to 199 mg dL, measured during the OGTT [10]. Although prediabetes is considered as an intermediate stage in the natural progression of T2DM [11], it has been reported that only 50 % of the subjects diagnosed with IGT developed diabetes within 10 years [12,13]. Moreover, long-term population studies have also shown that around 50 % of the diabetic patients did not exhibit IGT at any time prior to the diagnosis [14]. This suggests that the fasting and 2-hour plasma glucose levels used in and of themselves cannot accurately predict the future development of T2DM.

The availability of big data in the healthcare sector has made Machine learning (ML) a viable instrument for disease prediction [15,16]. In contrast to traditional diagnostic techniques employing population based statistics, ML methods develop models that are trained using large amounts of data. In a pilot study, Maeta et al developed a ML algorithm to predict the risk of developing glucose metabolism disorder using the OGTT data [17]. Barakat et al used socio-demographic information, and point-of-care testing from blood and urine to develop diagnostic models of diabetes [18]. This approach uses support vector machine (SVM) along with a rule-based explanation to provide a comprehensibility of the results to the clinicians. The plasma glucose levels at baseline and 2-hr were among the features used. Han et al employed an ensemble SVM and random forest learning approaches to develop a decision making algorithm for the diagnosis of diabetes [19]. However, investigations that are designed to identify individuals at high risk of developing T2DM in the long-term future are limited. The San Antonio diabetes prediction model (SADPM) [20] uses a logistic regression supported by physiological parameters such as systolic blood pressure and cholesterol level. The underlying causes of T2DM in the form insulin resistance and insulin secretion were studied to develop a prediction model in [14]. In another study, multivariate logistic models using the plasma glucose values measured in the OGTT were used to predict the future risk of developing T2DM [21,22]. The predictive power of different biomarkers such as the fasting plasma glucose level, BMI and hemoglobin A1C (HbA1c) for T2DM onset was assessed in [23]. This study focused on individuals with metabolic syndrome, a complex and serious health condition that greatly increases the risk of CVD and diabetes.

The standard ML algorithms are designed to yield optimal performance in terms of accuracy over the full dataset. However, medical applications such as diagnosis and prediction of a disease require a biased decision-making mechanism that favors one of the classes. This approach inherently maximizes the performance of the class that is more relevant in clinic terms. Therefore, the objective in such applications is to design a classifier that improves the accuracy of the class that is clinically more relevant. Additionally, often the amount of data is highly skewed with the clinically relevant class in an outsized minority. There are various roundabout ways to obtain accurate classifier performance in this scenario that include the method of sampling [24] in which the class distribution is artificially balanced by either under sampling the majority class, over-sampling the minority class or both. Furthermore, feature weighting schemes assign distinct costs to training examples [25] in order to introduce a certain bias. Other techniques introduce evaluation metric such as the geometric mean (g-mean) [26], that concurrently optimizes the positive class accuracy (sensitivity) and the negative class accuracy (specificity) [27].

We hypothesized that the features extracted from the OGTT will be able to predict the future onset of T2DM. In this paper, we therefore propose a screening tool that identifies the most relevant features extracted from the OGTT data that strongly correlate with the future development of T2DM. We then use SVM to develop a prediction model by utilizing these relevant features estimated from the longitudinal cohort study, the San Antonio Heart Study (SAHS) [28,29].

## Materials and methods

### San Antonio Heart Study

The SAHS is a population-based epidemiological study that was conducted to assess the risk factors of diabetes and cardiovascular diseases in healthy population [28,29]. In total, 5,158 men and non-pregnant women of Mexican American (MA) and Non-Hispanic White (NHW) residents of San Antonio, Texas participated in the study in two cohorts. The age of individuals at the time of recruitment was between 25 and 64 years. As a part of the data collection, plasma glucose and serum insulin concentrations were collected during the OGTT at the baseline and after an average follow-up of 7.5 years. The BMI was also recorded for each individual at the baseline. In this study, we analyzed only the data generated from the second cohort of the SAHS which comprised of 1,492 subjects from the second cohort of the SAHS.

T2DM was diagnosed at the follow-up using the WHO criteria, i.e. fasting glucose level >126 mg dL or 2-hr glucose level ≥200 mg dL [10]. Furthermore, all individuals taking anti-diabetic medications were also classified as having T2DM. Individuals that reported by themselves any cardiovascular event such as a heart attack, stroke or angina, were labeled as having CVD at the follow-up. All other participants without T2DM or self-reported CVD were labeled as healthy for the case of this study. During the course of this longitudinal study, a total of 171 individuals developed T2DM with 10 individuals also reporting at least one cardiovascular event. The incidence rate of T2DM in the second cohort of the SAHS population was 10.79 %. Table 1 shows the population distribution in terms of the four classes. The distribution in terms of the ethnicity shows the T2DM prevalence among the MA individuals more than double, as compared to the NHW population.

**Table 1.** The classification of the 1,492 subjects used in this study based on the ethnicity.

|  | Healthy | T2DM | CVD | T2DM+CVD |
| --- | --- | --- | --- | --- |
| Total | 1,277<br>85.56 % | 161<br>10.79 % | 44<br>2.95 % | 10<br>0.67 % |
| MA | 836<br>83.77 % | 131<br>13.13 % | 24<br>2.40 % | 7<br>0.70 % |
| NHW | 441<br>89.27 % | 30<br>6.07 % | 20<br>4.05 % | 3<br>0.61 % |

The data used in this study consists of plasma glucose and serum insulin concentrations sampled at the baseline, and at 30, 60 and 120 min thereafter. The individuals are labeled at the SAHS follow-up using the current standard of care [28]. Fig 1 shows the distributions of the data used in this study.

**Fig 1.**
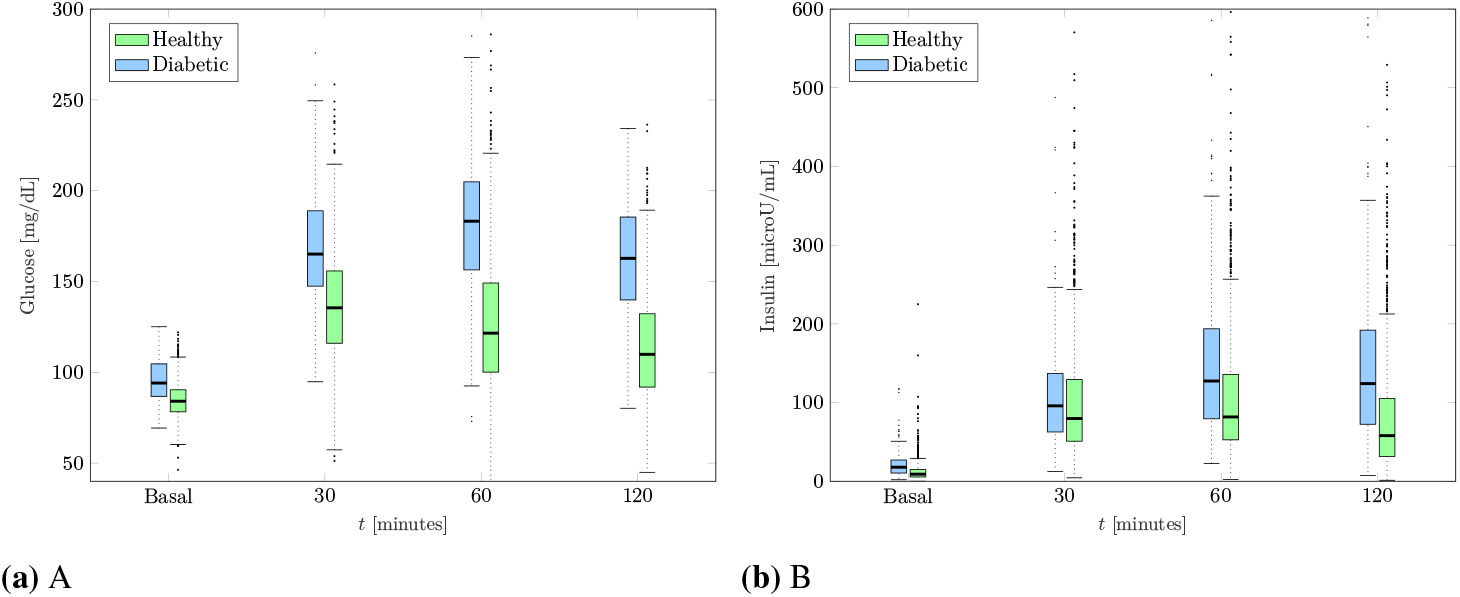
Box plots of glucose and insulin levels for healthy and diabetic subjects measured at the baseline OGTT. A: Plasma glucose. B: Serum insulin

### Machine Learning Framework

In this paper, we implemented SVM to construct the models for the prediction of future T2DM. The SVM develops models from a given training dataset such that it generalizes well to a new dataset and minimizes the empirical risk associated with misclassification of samples in the training set [30,31]. A model constructed by the SVM minimizes the overlap between classes in the training set by optimizing the separating hyperplane. For problems that may not be amenable to linear separation between the two classes, the SVM technique is very attractive due to the fact that the input feature space can be transformed to a higher dimension space, and a linear boundary can then be determined. This approach generally provides a better training performance, but potentially increases computational complexity excessively with the increase of the dimensionality of the input feature space [32]. The introduction of a kernel alleviates the need to determine the transformation by calculating the inner product between the coordinates of the input feature space instead. In this paper, we used the Gaussian radial basis function (RBF), as the kernel. The performance of the SVM can be optimized by tuning the free parameter of the kernel σ and specifying a cost that controls the rigidity of the class margin. This process is normally carried out through a grid search.

### Feature Extraction

We extracted all the features from the SAHS data acquired at the baseline. The dataset consists of plasma glucose and insulin concentrations recorded before glucose intake and at three time-points thereafter (30, 60, and 120 min). The labels (healthy and diabetes) were generated at the 7.5 years follow-up using the current standard of care diagnostics [28]. From the glucose and insulin concentrations, we computed the slope and area under the curve between all the possible combinations of a pair of measurements. In addition, we also calculated three empirical markers that describe the relationship between the glucose intake and insulin response. The first is the insulinogenic index (IGI) [33], which is a direct measure of the insulin response to glucose. It is calculated as the ratio of the slope of the insulin curve to the slope of the glucose curve between any two time intervals in the OGTT. The second marker, Matsuda index (M) evaluates the insulin sensitivity from the OGTT using a product of the weighted averages of the glucose and insulin concentrations [34],

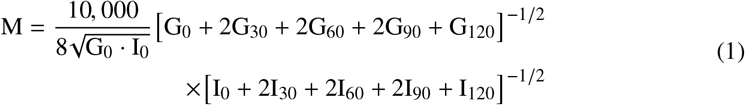

where the subscripts depict the time point of the OGTT. In case when the value at 90 min is not available, the average of 60 and 120 min is used instead [34]. The third marker, homeostatic model assessment - insulin resistance (HOMA-IR) [35] evaluates the beta-cell function. It is defined as the product of fasting plasma glucose concentration and fasting blood insulin concentration divided by 22.5. These markers have been used to estimate abnormalities in the insulin sensitivity. A total of 61 features (illustrated in Fig 2) are used in this study. The prefix AuC denotes the area under the curve and the slope is denoted by the symbol ∆. The term T_half_ represents the linearly interpolated value between any two intervals.

### Feature Selection

Before constructing the SVM model to predict a future diabetes occurrence, we search for the most effective subset of features in terms of relevance to the classifier output, i.e. incidence of T2DM at the follow-up. As a first step, we selected the ten most relevant features from the 61 available features using the minimum redundancy maximum relevance (mRMR) algorithm [36], which selects the most relevant features with minimum correlation among them. The mRMR algorithm determines the relevance between a feature (*x* as continuous random variable) and the class label (*y* as discrete random variable) in terms of the mutual information 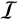 defined as [37],

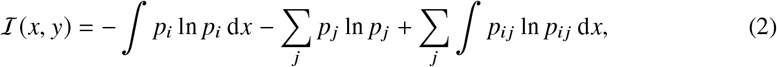

where *p*_*i*_, and *p*_*j*_ are the probabilities of the random variables *x* and *y* taking a particular value *x*_*i*_ and *y*_*j*_ ∈ (−1, 1) ∀ *j* respectively. The term *p*_*ij*_ denotes the joint probability *P*{*x* = *x*_*i*_, *y* = *y*_*j*_}. The three terms in Eq (2) represent the continuous, discrete and joint entropies of the random variables in the respective order. The features that are most relevant to the class label are the ones that maximize 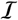. A heuristic approach is to keep only one a single feature from a correlated set of features that provides similar relevance information, and discard the remaining features. In order to ensure this, the mRMR algorithm minimizes the mutual correlation among the features expressed in terms of redundancy 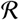,

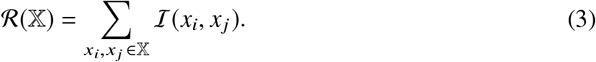

where 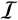 follows its definition in Eq (2). This procedure yielding maximum 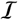 with respect to the diabetic class, along with minimal 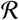, shortlists a set of ten features that are potentially strong predictors of the future development of T2DM.

**Fig 2.**
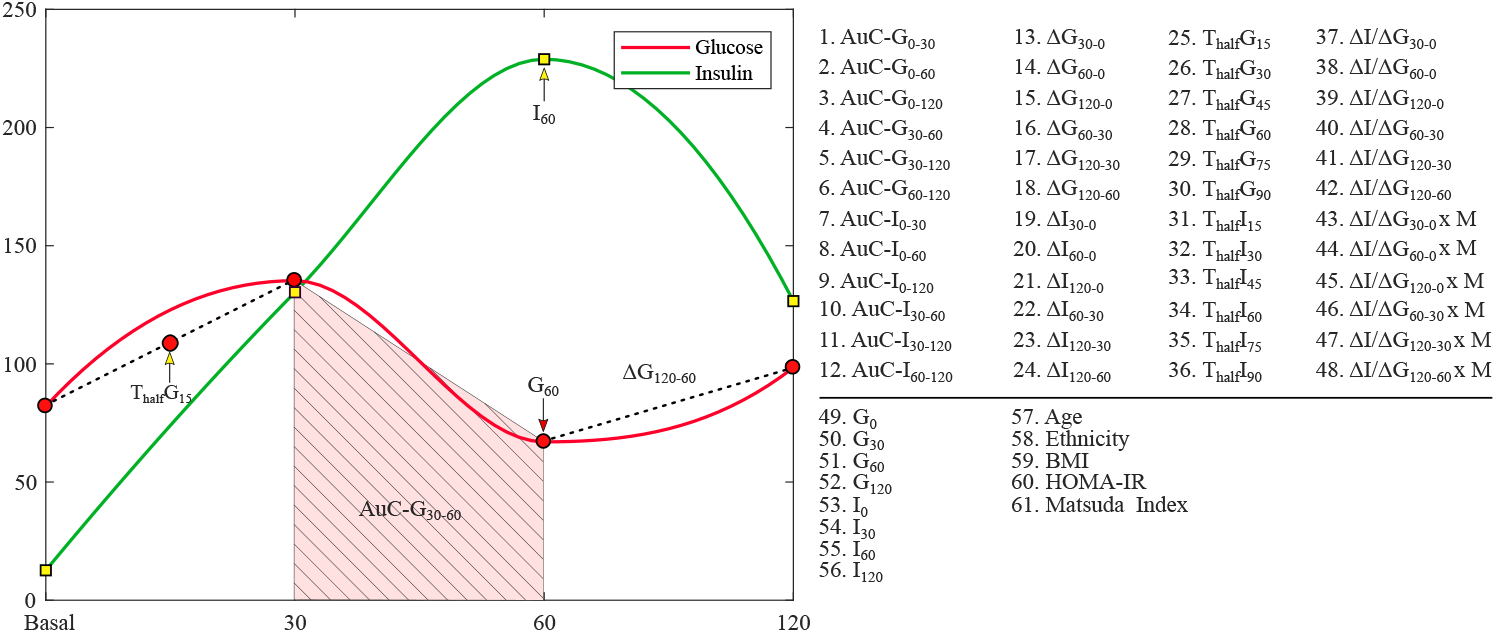
Illustration of all 61 features extracted from the SAHS dataset.

### Classification

We developed a supervised learning scheme using the baseline SAHS dataset and the labels (healthy, T2DM) obtained at the follow-up after an average of 7.5 years. In each experiment, we used a kernel-based binary SVM method to train, test and validate the performance of the diabetes prediction models. We excluded the 44 CVD entries as the only way of defining this class was based upon self-reporting and not on quantitative assessment. Furthermore, we also removed all entries with any information missing. That resulted in a total of 1,492 instances that were used in this study, out of which 171 were from the minority class and 1,321 were majority instances. As shown in Table 1, the SAHS dataset is intrinsically unbalanced with the class distribution skewed toward the majority class with a ratio of 7.5:1. We considered the minority class of diabetic subjects as the positive class with a label of 1, whereas the majority class consisting of healthy persons was termed as the negative class marked by a ‘-1’ label. To standardize the feature range prior to training, the feature space was scaled to unit variance around the respective mean for each feature respectively. To ensure that a model was unbiased, robust, and generalized well to the new data, we performed 10-fold cross-validation (CV).

For each CV, we first randomly selected a hold-out set consisting of 11 minority and 83 majority instances. We evaluated each model 100 times, in which the data was randomly partitioned on each occasion. We compared the performances of linear and non-linear SVM for all 1,023 possible combinations of the 10 most relevant features by considering all 1 to 10 combinations of features?. The optimal hyperplane parameters of the kernel were determined through a grid search. To select the best feature set, we have used the geometric mean of sensitivity and specificity [26]. All experiments were performed by an in-house developed software using Matlab®(version 9.2.0 MathWorks Inc., Natick, Massachusetts, USA).

## 1 Results and Discussion

The mRMR algorithm produces a sequential list of ten ranked features, shown in Table 2. Besides ethnicity (ranked fourth), all other features are notably derived from OGTT measurements. The list contains six features derived from plasma glucose concentrations, while only three features are deduced from insulin concentrations.

**Table 2.** List of ten most relevant features ranked by the mRMR algorithm

| Rank | Feature |
| --- | --- |
| 1 | AuC-G<sub>0-120</sub> |
| 2 | $\Delta G_{120-0}$ |
| 3 | $\Delta G_{120-60}$ |
| 4 | Ethnicity |
| 5 | $\Delta I_{120-0}$ |
| 6 | $\Delta G_{60-0}$ |
| 7 | $\Delta G_{30-0}$ |
| 8 | $\Delta G_{60-30}$ |
| 9 | $\Delta I_{120-60}$ |
| 10 | $\Delta I_{60-0}$ |

In all the classification experiments, we aimed to maximize the ability to correctly predict the diabetic class without compromising the classifier accuracy. The bar plots in Fig. 3 show the g-mean of the sensitivity and specificity obtained from the linear and RBF kernels. For each number of features used, we selected the combination that generated the maximum g-mean. All the results presented here are averaged over 100 iterations of the respective classifiers. The g-mean obtained from the linear SVM ranges from 0.8711 to 0.8742. As observed from Fig. 3a, the addition of more features does not result in a substantial performance improvement. However, the maximum g-mean of the sensitivity and specificity is obtained when all 10 features are used. For the non-linear SVM with RBF kernel, the g-mean ranges from 0.8638 to 0.8903. The combination of the features namely, AuC-Glu_0-120_, ∆G_120-0_, ∆G_120-60_ and ∆G_30-0_ yields the maximum performance. Notably, all four features are derived from the plasma glucose concentrations. We note that the glucose derived features are ranked the highest during feature selection. Moreover, a combination of glucose only features generate the best SVM models when less than four features are used.

**Fig 3.**
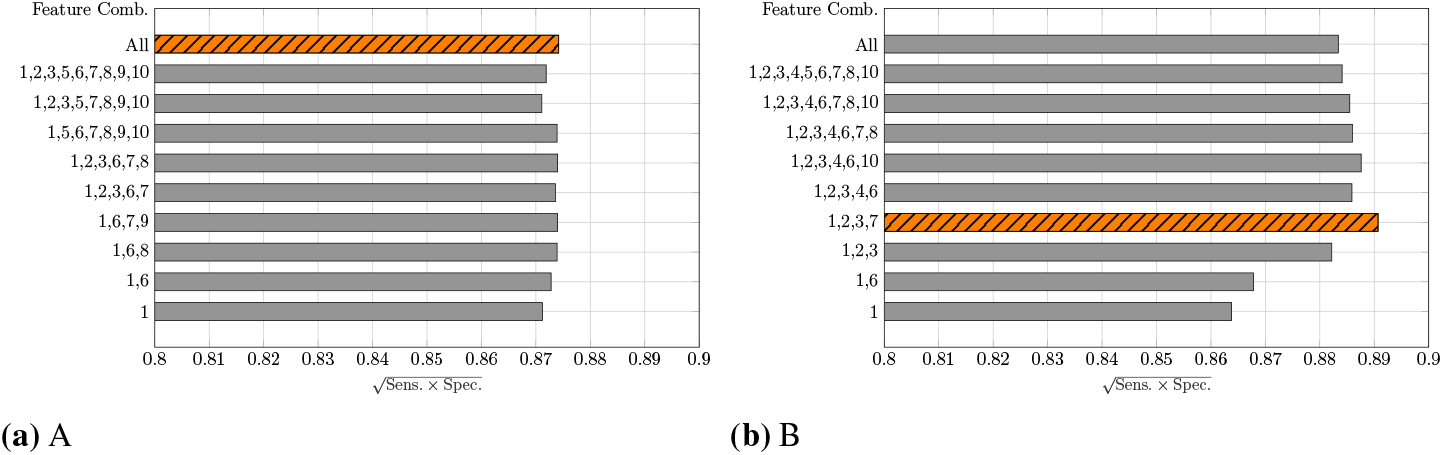
The g-mean of sensitivity and specificity for A: linear, and B: RBF kernels. The maximum performance feature combination is depicted by a different color scheme.

The accuracy and sensitivity of the same feature combinations are separately shown in Fig. 4. The best model obtained using a combination of four glucose derived features and RBF kernel has an accuracy of 96.80 %, and sensitivity of 80.09 %.

**Fig 4.**
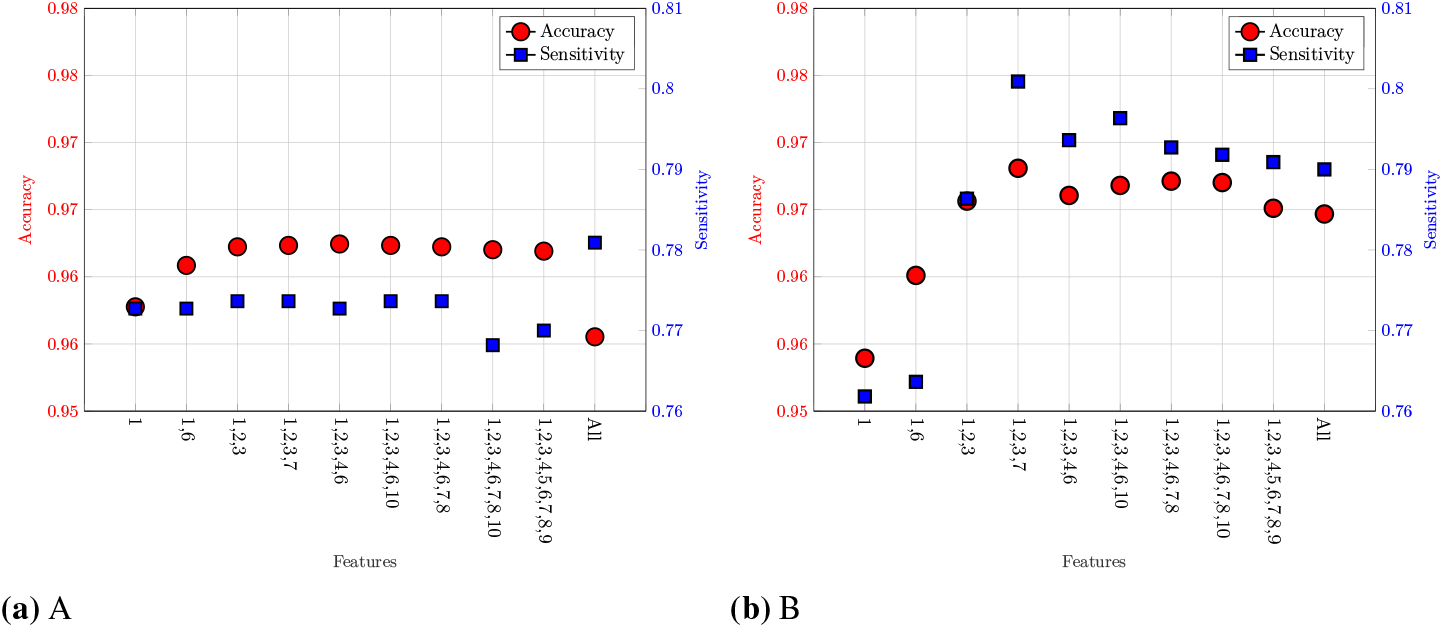
The classifier performance in terms of accuracy and sensitivity for the best feature combinations. A: Linear kernel. B: RBF kernel.

Table 3 presents a comparison of the generated SVM models to the results obtained in other studies using the SAHS dataset. We compared our results with the SADPM [20], in which a person’s age, gender, ethnicity, fasting glucose level, family history, blood pressure, and cholesterol level were used to construct a logistic regression. It is notable that the SADPM has the highest sensitivity (88.80 %) however, the increased prediction performance comes along with a very low accuracy of 56.33 %. In [21], a two-step approach was introduced that first used the SADPM risk score and then augmented it with the 1-hour plasma glucose concentration measured in the OGTT. This strategy resulted in an improved accuracy but the sensitivity dropped to 77.70 %.

**Table 3.** Comparison of validation performance of the best SVM classifiers with previous studies.

| | Accuracy ± SD | Specificity ± SD | Specificity ± SD | g-mean ± SD |
| --- | --- | --- | --- | --- |
| Linear SVM (10 features) | 95.55 % ± 0.24 % | 78.09 % ± 0.33 % | 97.87 % ± 0.30 % | 0.8742 ± 0.2100 |
| SVM-RBF (4 features) | 96.80 % ± 0.41 % | 80.09 % ± 1.42 % | 99.02 % ± 0.33 % | **0.8903 ± 1.5600** |
| SADPM [20] | 56.329 % | 88.80 % | 52.00 % | 0.6795 |
| Two-step Approach [21] | 77.43 % | 77.70 % | 77.40 % | 0.7755 |

In the SAHS dataset, the prevalence of IFG and IGT was 8.91 % (133 instances) and 22.52 % (336 instances) respectively. Out of the 399 subjects diagnosed with prediabetes showing IFG or IGT at the baseline, only 120 (30.08 %) actually developed diabetes between the baseline and the follow-up. Furthermore, 120 (25.67 %) subjects diagnosed with diabetes at the follow-up did not show any symptoms of either IGT or IFG at the baseline.

Our investigation shows that features derived from insulin have less predictive value for T2DM as compared to glucose based features. Indices such as Matsuda and HOMA-IR that are commonly used to assess the insulin function, also did not yield high correlation with the future development of T2DM.

## 2 Conclusion

In this paper, we present a most promising set of features that are used to develop a non-linear SVM based future T2DM prediction model. The features were derived from the OGTT data and were augmented by personal information such as age, ethnicity, and BMI. Using a feature selection algorithm, we demonstrate that the features deduced from the plasma glucose concentrations provide the optimal feature subset and have the strongest predictive power for the future development of T2DM. Moreover, the performance of the presented prediction model is significantly better in terms of combined accuracy and sensitivity combined, compared to other T2DM prediction models. In order to address the unbalanced nature of the SAHS dataset, we chose the g-mean of sensitivity and specificity as the performance evaluation criteria.

The principal contribution of this study includes a T2DM prediction model based on the features derived only from the plasma glucose concentrations measured during an OGTT. The findings of this paper provide a complementary and cost-effective tool for the clinicians to screen individuals that are at an increased risk of developing T2DM in the future.

## Acknowledgments

This publication was made possible by NPRP grant number NPRP 10-1231-160071 from the Qatar National Research Fund (a member of Qatar Foundation). The statements made herein are solely the responsibility of the authors.

## References

1. Mathers CD, Loncar D. Projections of Global Mortality and Burden of Disease from 2002 to 2030. PLoS Medicine. 2006;3(11):e442.

2. Tuomilehto J, Lindström J, Eriksson JG, Valle TT, Hämäläinen H, Ilanne-Parikka P, et al. Prevention of type 2 diabetes mellitus by changes in lifestyle among subjects with impaired glucose tolerance. New England Journal of Medicine. 2001;344(18):1343–1350.

3. Diabetes Prevention Program Research Group. Long-term effects of lifestyle intervention or metformin on diabetes development and microvascular complications over 15-year follow-up: the Diabetes Prevention Program Outcomes Study. The Lancet Diabetes & Endocrinology. 2015;3(11):866–875.

4. Noble D, Mathur R, Dent T, Meads C, Greenhalgh T. Risk models and scores for type 2 diabetes: systematic review. BMJ. 2011;343:d7163.

5. Heikes KE, Eddy DM, Arondekar B, Schlessinger L. Diabetes Risk Calculator. Diabetes Care. 2008;31(5):1040–1045.

6. Glümer C, Carstensen B, Sandbæk A, Lauritzen T, Jørgensen T, Borch-Johnsen K. A Danish Diabetes Risk Score for Targeted Screening. Diabetes Care. 2004;27(3):727–733.

7. Heliövaara M, Aromaa A, Klaukka T, Knekt P, Joukamaa M, Impivaara O. Reliability and validity of interview data on chronic diseases The mini-Finland health survey. Journal of Clinical Epidemiology. 1993;46(2):181–191.

8. Expert Committee on the Diagnosis and Classification of Diabetes Mellitus. Report of the Expert Committee on the Diagnosis and Classification of Diabetes Mellitus. Diabetes Care. 1997;20(7):1183–1197.

9. Stumvoll M, Mitrakou A, Pimenta W, Jenssen T, Yki-Järvinen H, Van Haeften T, et al. Use of the oral glucose tolerance test to assess insulin release and insulin sensitivity. Diabetes Care. 2000;23(3):295–301.

10. World Health Organization, International Diabetes Federation. Definition and diagnosis of diabetes mellitus and intermediate hyperglycaemia: report of a WHO/IDF consultation. World Health Organization; 2006.

11. DeFronzo RA, Abdul-Ghani M. Assessment and treatment of cardiovascular risk in prediabetes: Impaired glucose tolerance and impaired fasting glucose. The American Journal of Cardiology. 2011;108(3):3B–24B.

12. Shaw JE, Zimmet PZ, de Courten M, Dowse GK, Chitson P, Gareeboo H, et al. Impaired fasting glucose or impaired glucose tolerance. What best predicts future diabetes in Mauritius? Diabetes Care. 1999;22(3):399–402.

13. Unwin N, Shaw J, Zimmet P, Alberti KGMM. Impaired glucose tolerance and impaired fasting glycaemia: the current status on definition and intervention. Diabetic Medicine. 2002;19(9):708–723.

14. Abdul-Ghani MA, Williams K, DeFronzo RA, Stern M. What Is the Best Predictor of Future Type 2 Diabetes? Diabetes Care. 2007;30(6):1544–1548.

15. Erraguntla M, Zapletal J, Lawley M. Framework for Infectious Disease Analysis: A comprehensive and integrative multi-modeling approach to disease prediction and management. Health Informatics Journal. 2017; p. 1460458217747112.

16. Freeze J, Erraguntla M, Verma A. Data Integration and Predictive Analysis System for Disease Prophylaxis: Incorporating Dengue Fever Forecasts. In: Proceedings of the Hawaii International Conference on System Sciences (HICSS); 2018. p. 1–10.

17. Maeta K, Nishiyama Y, Fujibayashi K, Gunji T, Sasabe N, Iijima K, et al. Prediction of Glucose Metabolism Disorder Risk Using a Machine Learning Algorithm: Pilot Study. JMIR Diabetes. 2018;3(4):e10212. doi:10.2196/10212.

18. Barakat N, Bradley AP, Barakat MNH. Intelligible Support Vector Machines for Diagnosis of Diabetes Mellitus. IEEE Transactions on Information Technology in Biomedicine. 2010;14(4):1114–1120.

19. Han L, Luo S, Yu J, Pan L, Chen S. Rule Extraction From Support Vector Machines Using Ensemble Learning Approach: An Application for Diagnosis of Diabetes. IEEE Journal of Biomedical and Health Informatics. 2015;19(2):728–734.

20. Stern MP, Williams K, Haffner SM. Identification of persons at high risk for type 2 diabetes mellitus: do we need the oral glucose tolerance test? Annals of Internal Medicine. 2002;136(8):575–581.

21. Abdul-Ghani MA, Abdul-Ghani T, Stern MP, Karavic J, Tuomi T, Bo I, et al. Two-Step Approach for the Prediction of Future Type 2 Diabetes Risk. Diabetes Care. 2011;34(9):2108–2112.

22. Abdul-Ghani MA, Lyssenko V, Tuomi T, DeFronzo RA, Groop L. Fasting versus postload plasma glucose concentration and the risk for future type 2 diabetes: results from the Botnia Study. Diabetes Care. 2009;32(2):281–286.

23. Ozery-Flato M, Parush N, El-Hay T, Visockienė Ž, Ryliškytė L, Badarienė J, et al. Predictive models for type 2 diabetes onset in middle-aged subjects with the metabolic syndrome. Diabetology & Metabolic Syndrome. 2013;5(1):36.

24. Chawla NV, Bowyer KW, Hall LO, Kegelmeyer WP. SMOTE: synthetic minority over-sampling technique. Journal of Artificial Intelligence Research. 2002;16:321–357.

25. Domingos P. MetaCost: A General Method for Making Classifiers Cost-sensitive. In: Proceedings of the Fifth ACM SIGKDD International Conference on Knowledge Discovery and Data Mining. KDD ’99. New York, NY, USA: ACM; 1999. p. 155–164.

26. Kubat M, Matwin S, et al. Addressing the curse of imbalanced training sets: one-sided selection. In: ICML. vol. 97. Nashville, USA; 1997. p. 179–186.

27. Tang Y, Zhang Y, Chawla NV, Krasser S. SVMs Modeling for Highly Imbalanced Classification. IEEE Transactions on Systems, Man, and Cybernetics, Part B (Cybernetics). 2009;39(1):281–288.

28. Burke JP, Williams K, Gaskill SP, Hazuda HP, Haffner SM, Stern MP. Rapid Rise in the Incidence of Type 2 Diabetes From 1987 to 1996: Results From the San Antonio Heart Study. Archives of Internal Medicine. 1999;159(13):1450.

29. Lorenzo C, Williams K, Hunt KJ, Haffner SM. Trend in the Prevalence of the Metabolic Syndrome and Its Impact on Cardiovascular Disease Incidence: The San Antonio Heart Study. Diabetes Care. 2006;29(3):625–630.

30. Vapnik VN. The nature of statistical learning theory. 2nd ed. Statistics for engineering and information science. New York: Springer; 2000.

31. Vapnik VN, Chervonenkis AY. On the uniform convergence of relative frequencies of events to their probabilities. In: Measures of complexity. Springer; 2015. p. 11–30.

32. Friedman J, Hastie T, Tibshirani R. The elements of statistical learning. Springer Series in Statistics. Springer New York; 2001.

33. Seino Y, Ikeda M, Yawata M, Imura H. The insulinogenic index in secondary diabetes. Hormone and Metabolic Research. 1975;7(02):107–115.

34. Matsuda M, DeFronzo RA. Insulin sensitivity indices obtained from oral glucose tolerance testing: comparison with the euglycemic insulin clamp. Diabetes Care. 1999;22(9):1462–1470.

35. Matthews D, Hosker J, Rudenski A, Naylor B, Treacher D, Turner R. Homeostasis model assessment: insulin resistance and β-cell function from fasting plasma glucose and insulin concentrations in man. Diabetologia. 1985;28(7):412–419.

36. Peng H, Long F, Ding C. Feature selection based on mutual information: criteria of max-dependency, max-relevance, and min-redundancy. IEEE Transactions on Pattern Analysis and Machine Intelligence. 2005; p. 1226–1238.

37. Ross BC. Mutual information between discrete and continuous data sets. PloS one. 2014;9(2):e87357.

